# Strategies and Techniques for Quality Control and Semantic Enrichment with Multimodal Data: A Case Study in Colorectal Cancer with eHDPrep

**DOI:** 10.1101/2022.09.07.506953

**Authors:** Tom M Toner, Paul Miller, Thorsten Forster, Helen G Coleman, Ian M Overton

**Affiliations:** Patrick G Johnston Centre for Cancer Research, Queen’s University Belfast, Belfast, UK; Health Data Research Wales and Northern Ireland, Queen’s University Belfast, Belfast, UK; The Centre for Secure Information Technologies, Queen’s University Belfast, UK; LifeArc, Nine, Edinburgh BioQuarter, 9 Little France Road, Edinburgh, UK; Centre for Public Health, Queen’s University Belfast, Belfast, UK

**Keywords:** Quality Control, Semantic Enrichment, Ontology, Colorectal Cancer, Health data, Medical Informatics, Quality Assessment, Data integration, Bioinformatics

## Abstract

**Background:** Integration of data from multiple domains can greatly enhance the quality and applicability of knowledge generated in analysis workflows. However, working with health data is challenging, requiring careful preparation in order to support meaningful interpretation and robust results. Ontologies encapsulate relationships between variables that can enrich the semantic content of health datasets to enhance interpretability and inform downstream analyses.

**Findings:** We developed an R package for electronic Health Data preparation ‘eHDPrep’, demonstrated upon a multi-modal colorectal cancer dataset (n=661 patients, n=155 variables; Colo-661). eHDPrep offers user-friendly methods for quality control, including internal consistency checking and redundancy removal with information-theoretic variable merging. Semantic enrichment functionality is provided, enabling generation of new informative ‘meta-variables’ according to ontological common ancestry between variables, demonstrated with SNOMED CT and the Gene Ontology in the current study. eHDPrep also facilitates numerical encoding, variable extraction from free-text, completeness analysis and user review of modifications to the dataset.

**Conclusion:** eHDPrep provides effective tools to assess and enhance data quality, laying the foundation for robust performance and interpretability in downstream analyses. Application to a multi-modal colorectal cancer dataset resulted in improved data quality, structuring, and robust encoding, as well as enhanced semantic information. We make eHDPrep available as an R package from CRAN [[URL will go here]].

## Background

Health data can be challenging to work with, arising from incompleteness, fragmentation, inaccuracies and the presence of unstructured information [1]. Data quality is an essential parameter for productive analysis, widely recognised in the adage ‘garbage in - garbage out’ [2]. Thus, quality control (QC) procedures, including quality assessment, lay foundations for drawing robust conclusions from health data. The fundamental dimensions of data quality are consistency, accuracy, completeness, record uniqueness, timeliness and validity (syntactic conformity) [3,4]. Applicability is a further important consideration for data quality; encoding data in a numeric and machine interpretable format is vital for accurate interpretation in advanced analysis workflows [5]. Ontologies provide structured representations of a knowledge domain and can support QC when dataset variables are mapped to ontological entities. For instance, multiple variables may map to the same or semantically similar concepts, suggesting opportunities for merging operations or internal consistency checks [6]. Ontologies also provide computable information on the semantic relationships between terms which can add value to downstream analysis [7]. The semantic information held in ontologies can be leveraged to generate new variables through aggregation of existing variables during post-QC data preparation in a process we describe as semantic enrichment.

Several tools are available for health data QC, however these are typically aimed at single modalities. For example, ‘dataquieR’ (completeness, consistency, accuracy, validity) and ‘mosaicQA’ (completeness, validity) focus upon observational health and epidemiological research data [8,9]. Packages such as ‘summarytools’ offer more generalised functionality to facilitate data exploration through summary descriptive reports (completeness, accuracy) [10]. Other packages support targeted encoding such as ‘genetics’ which targets genetic data (i.e. genotypes and haplotypes) [11] while ‘quanteda’ provides extensive tools for natural language processing [12]. The ‘tidyverse’ collection builds upon base R’s functionality to improve the capability, efficiency, and programmability of data scientists’ QC workflows [13,14]. Several R packages calculate semantic similarities [15–17] however we are not aware of any which provide the ability to aggregate variables using semantic commonalities in preparation for analysis.

QC may require up to approximately 80% of a data mining project’s time [18]. While data quality and encoding issues in multimodal data can currently be tackled by combining multiple existing approaches, each requires time-consuming familiarisation and may require multiple data transformations potentially adversely impacting data quality [4]. We present a toolkit for electronic Health Data Preparation (eHDPrep), enabling robust programmatic QC and enrichment of semantic content; high-level functions empower general R users to assess, process, and review their dataset with minimal coding while low-level functions allow advanced Rusers to specify parameters and workflows as required. We demonstrate the utility of eHDPrep on a multimodal dataset containing 155 variables for 661 colon cancer patients (Colo-661) [19,20]. Colorectal cancer (CRC) has a large disease burden as the third most common malignancy with an estimated 1.9 million new cases and 915,800 deaths worldwide in 2020 [21]; advances in CRC medicine are urgently needed [22].

## Findings

### Quality Control

Data reliability encompasses completeness, consistency, accuracy, uniqueness, and validity [3,4]; eHDPrep addresses issues in these dimensions through both specific low-level functions and in the high-level functions ‘assess_quality’, ‘apply_quality_ctrl’, and ‘review_quality_ctrl’. The QC workflow in eHDPrep provides user-friendly methods to evaluate and address data quality issues (Figure 1). We present the application of this workflow to Colo-661 in the sections below in order to enhance data reliability, to enable machine interpretability, and to assess the effects of QC operations upon the dataset.

**Figure 1:**
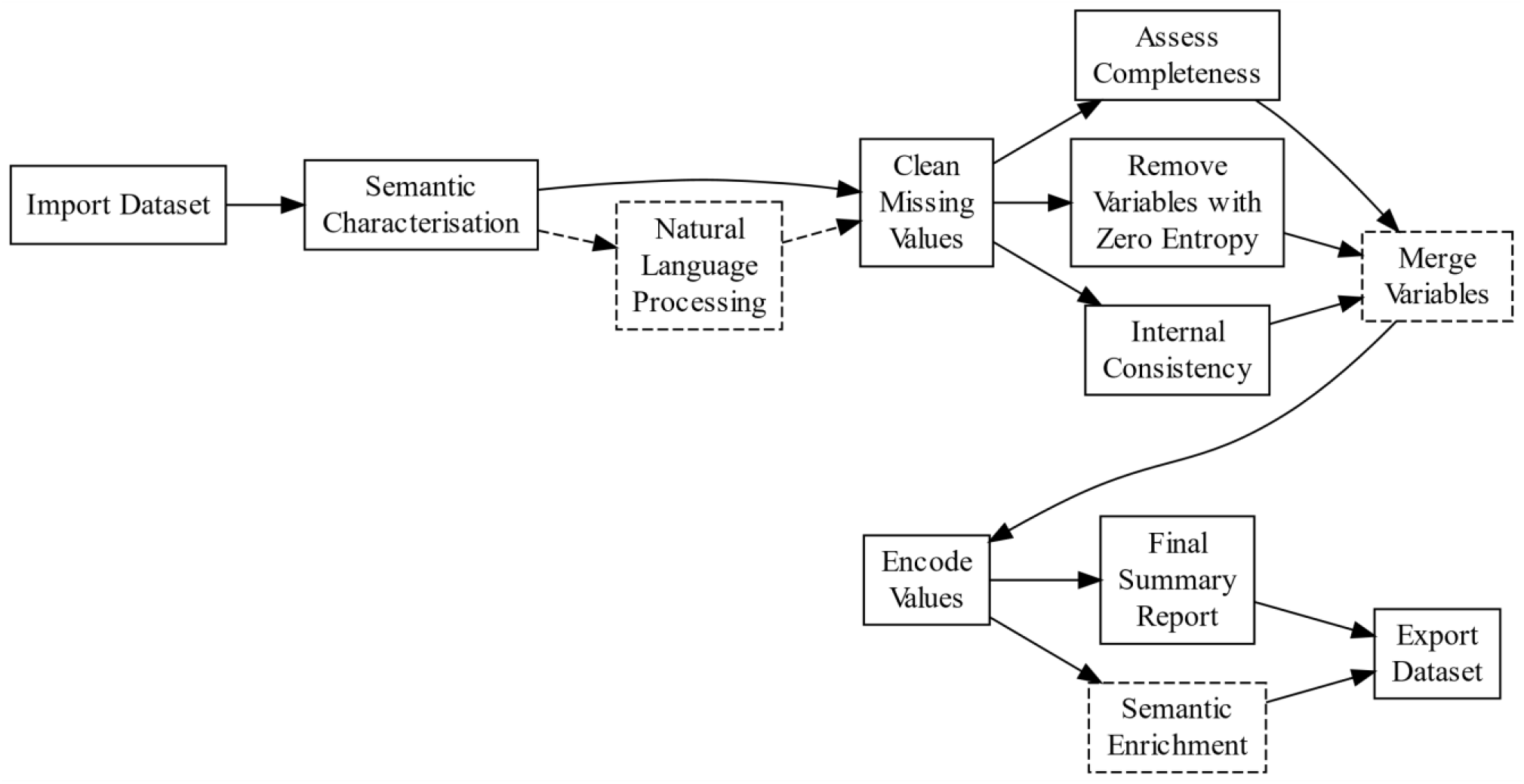
Overview of eHDPrep workflow. The ordering of steps reflects logical dependencies. Dashed arrows and boxes signify optional steps. Following import, the semantic characteristics of the data are established, missing values are dealt with and a series of operations may be performed. ‘Natural Language Processing’ is only required if free-text variables are present. ‘Merge Variables’ is an optional step for user-defined merging operations with functionality to measure information loss. Variables are encoded in a machine interpretable format and a summary report is generated for review by the user. Additionally, functionality to review each step is provided. ‘Semantic Enrichment’ optionally involves aggregation of variables according to semantic commonalities identified by an ontology such as SNOMED CT [23].

### Semantic Characterisation

Following data import, semantic characterisation is required in order to determine variable classifications (Supplementary Table 1) along with information provided by the user, for example regarding data modalities. The semantic characterisation process includes user review of the automated variable type assignments. Correct semantic information is essential for successful application of downstream steps in eHDPrep.

### Natural Language Processing

Analysis workflows typically require data in a standardised format, however significant health data are contained within free-text clinical notes [24]. eHDPrep includes user-friendly extraction of information from free-text by wrapping Natural Language Processing functionality from quanteda [12] and tm [25] to create variables describing frequently occurring words or groups of nearby words (eHDPrep function ‘extract_freetext’). Three free-text variables in Colo-661, containing digitised medical notes, were transformed into eleven new structured variables. Of these, six variables were generated from family members’ cancer history (recorded in 21% of patients) following manual correction of observed misspellings, expansion of abbreviations, and standardising cancer name to ‘[cancer location] cancer’ (e.g. “melanoma” to “skin cancer”). Four of the new structured variables identified the occurrence of cancer in close family members (mother, brother, sister, father), two further variables recorded if a family member had lung or breast cancer. Manual review determined that the data extraction for these variables had 89.9% sensitivity and 99.9% specificity across the generated values. False positives and negatives were manually corrected in Colo-661.

### Encoding Missing Values

Proper representation of missing values is critically important for the correct execution of downstream functions, for example if missing values are to be excluded from calculations. Missing values may be encoded in a variety of ways, including strings (e.g. ‘missing’, ‘unknown’) or out-of-range values (e.g. ‘-1’) [26]. Indeed, missing values in Colo-661 were recorded in eight encodings, representing 4.3% of dataset values, which were converted to ‘NA’ values using eHDPrep.

### Completeness

The degree to which a dataset is populated with data, rather than missing values, is a vital early measurement in quality assessment. eHDPrep measures both variable and patient record completeness at a whole-dataset scale, visualised across Colo-661 in Supplementary Figure S1. Patterns of completeness may also be explored with eHDPrep through a binary heatmap; the clusters of missing data in Colo-661 showed good correspondence with different data types demonstrating non-random missingness (Figure 4a). Variables with zero entropy [27] (Equation (1) have the same value across all records and, for example, cannot be used to stratify the cohort. Zero entropy variables therefore have limited utility, even if fully complete, and are flagged by eHDPrep. Four Colo-661 variables were removed due to zero entropy. These quality assessment procedures are achieved using the functions ‘assess_completeness’ and ‘assess_quality’.

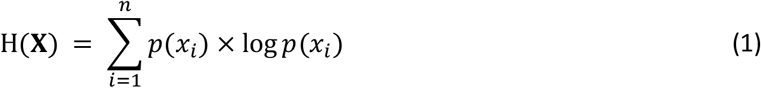

Where *p*(*x*_*i*_) is the probability of each element *x* occurring in the input vector **X**.

### Internal Consistency

In order to enable evaluation of internal inconsistencies, eHDPrep assesses user-supplied semantic dependencies between variable pairs. In such dependencies, a value in one variable limits the logically valid values in the other. We designed forty-nine internal consistency checks for Colo-661 across fifteen variables (with some variables present in multiple pairs; Supplementary Table S2). As expected in real-world data, we found forty instances of internal inconsistency across five variable pairs, demonstrating the value of this automated approach.

There was a conflict between the related variables ‘N stage’ and ‘number of positive lymph nodes’ (Figure 2a). One record had a value of ‘N2’ for the ‘N stage’ variable; however, the ‘number of positive lymph nodes’ value was lower than required for assignment of N2 status according to the staging criteria [28]. Similarly, we identified three records where the ‘number of lymph nodes examined’ was fewer than the ‘number of positive lymph nodes’ (Figure 2b). Thirty records contained inconsistencies due to a category mislabelling in relation to tumour budding [29] (‘high, >10’ instead of ‘high, >=10’) which was identified when comparing a discretized variable with its corresponding non-discretized variable. Four records stated that patients did not have a personal history of cancer while stating that the patient had non-melanoma skin cancer. Two records stated that the patient had a hereditary form of CRC while stating that the patient had no or an unknown family history of CRC.

**Figure 2:**
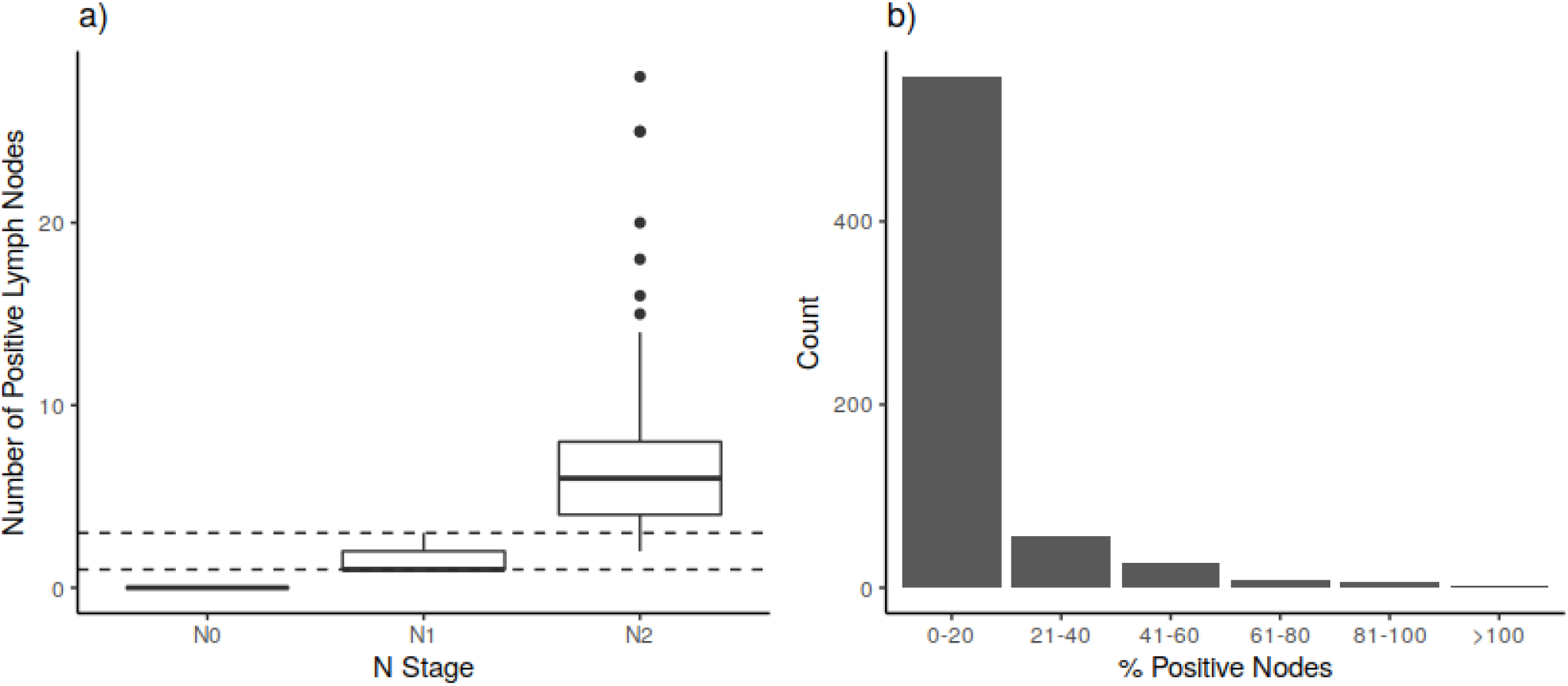
Automated identification of internal inconsistencies with eHDPrep. a) A box plot comparing values for ‘number of positive lymph nodes’ (y-axis) and ‘N stage’ (x-axis) in Colo-661. Boxes span the interquartile range with an internal line for the median; whiskers represent the maximum value of the data within 1.5 times the interquartile range beyond the 75th or 25th percentile for the upper or lower whiskers, respectively. Dashed lines show bounds on the ‘number of positive lymph nodes’ for classification into the ‘N Stage’ category N1. A portion of records are inconsistent because they have ‘N stage’ value N2 while the ‘number of positive lymph nodes’ indicates ‘N stage’ of N1. b) The percentage of positive lymph nodes is shown, derived from analysis of records for the ‘number of positive lymph nodes’ against the ‘number of lymph nodes examined’. Where values exceed 100% (n=3) there is a logical inconsistency, because the ‘number of positive lymph nodes’ should not exceed the ‘number of lymph nodes examined’.

Flagging the above inconsistencies focussed further data curation in order to resolve these conflicts. In the above instances we removed any inconsistent values from one variable in the pair, selected by assessing the reliability of the data source. A more conservative strategy might be required if expert curation is not possible; for example, involving elimination of all conflicting values or potentially removing the inconsistent variables entirely.

### Variable Merging

Merging variables can improve analysis by reducing redundancy and improving storage efficiency. However, inappropriate merging may lead to information loss. Accordingly, we developed functionality in eHDPrep for quantitative evaluation of merging operations using an information theoretic approach. Information Content (IC; Equation 2) is determined from category probabilities for discrete variables, or with variable bandwidth kernel density estimation for continuous variables [30]. The Mutual Information Content (MIC; Equation 3) of each input variable with the merged variable is also calculated [30,31]. Potential information loss during variable merging can be assessed by comparing the MIC of an input variable and the merged variable against the IC of the input variable. If the MIC and IC are identical, the input variable’s information is retained within the merged variable.

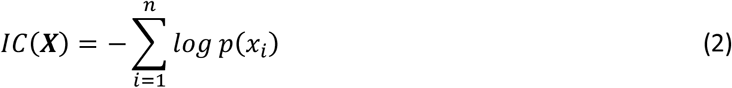

Where *p*(*x*_*i*_) is the probability of each element *x* occurring in the input vector ***X***.

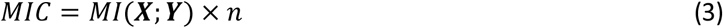

Where ***X*** and ***Y*** are numeric vectors, *MI* is mutual information, and *n* is the number of complete cases in both ***X*** and ***Y***.

As an example of support for variable merging in eHDPrep, Figure 3 visualises two candidate merging operations applied to Colo-661 variables describing a scoring of Crohn’s-like lymphoid reaction in the tumour [32] based on Graham-Appelman criteria [33]. One possible merging strategy aggregates values ‘1’ and ‘2’ from Input 1 to ‘1-2’ in the merged variable, leading to information loss (Figure 3a). A superior strategy takes the values ‘1-2’ from Input 2 as an ordinal category value between ‘1’ and ‘2’ and does not produce any information loss (Figure 3b).

**Figure 3:**
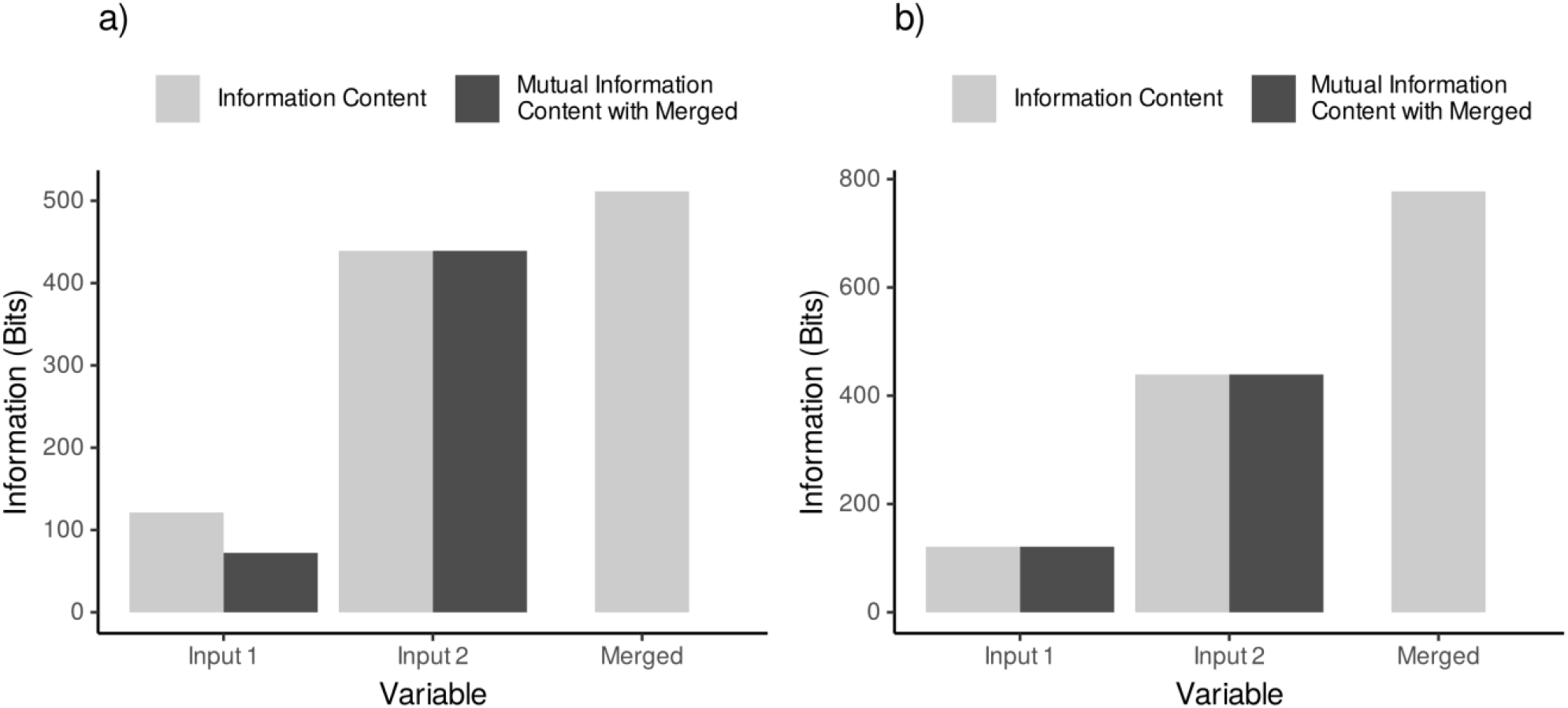
Information theoretic evaluation of merging operations. A comparison of two candidate merging approaches for variables pertaining to Crohn’s-like lymphoid reaction in the tumour. a) Merging through aggregation of the Input 1 values ‘1’ and ‘2’ to ‘1-2’ to align with Input 2’s ‘1-2’ value. The Mutual Information Content (MIC) of Input 2 with the merged variable is equal to the information content (IC) of Input 2; hence all of the information from Input 2 is captured by the merged variable. In contrast, Input 1 has IC higher than its MIC with the merged variable and so information loss has occurred. b) Lossless merging operation where ‘1-2’ values of Input 2 were encoded as an intermediate ordinal category between ‘1’ and ‘2’ from Input 1. All information from both input variables is contained in the merged variable (i.e. IC is equal to MIC with the merged variable). Indeed, the IC of the merged variable in b) is greater than the value shown in a). Therefore, the merging operation shown in b) is advantageous.

### Encode Values

A total of 123 structured variables in Colo-661 were numerically encoded for enhanced machine interpretability. Twenty-two ordinal and sixty-nine binary category variables were encoded as ordered factors, allowing numerical representation of ordinal relationships between values while preserving the original labelling. For example, tumour N stages [28] N0, N1, N2 were encoded as 1, 2, 3. Nineteen variables measuring single substitution mutations were also encoded as ordered factors where the order was determined by the frequency of the mutation status in the cohort; the most common status was encoded as ‘0’ and least common status encoded as ‘2’. Thirteen non-binary nominal variables were transformed into binary variables describing the presence of each unique value in the source variable using one-hot encoding (Supplementary Table S1) [5]. Following the above encoding steps, human-interpretable labels in ordinal variables were transferred to a mapping reference table at the end of QC during an assertion confirming that all variables were numeric following QC, in contrast to 16.8% before processing with eHDPrep.

### Quality Review

Understanding the effect of QC operations applied across large health datasets is non-trivial; eHDPrep simplifies this process and concisely records data changes resulting from QC. Firstly, eHDPrep can produce a comparative tally of unique combinations of values in variables before and after a change was implemented. These tallies can be produced after each QC action, showing the incremental changes to the dataset. Secondly, eHDPrep facilitates final review of changes to variable count which measured 207 variables in Colo-661 following QC, or value-level QC modifications which are optionally summarised in a bar plot. This plot (Figure 5) highlights differences in the proportion of values modified across Colo-661 that may inform upon the underlying structure of the dataset. Thirdly, eHDPrep’s comparative completeness function visualises the distribution of variable or row completeness before and after QC. Figure 4b demonstrates the positive impact of eHDPrep QC on Colo-661’s variable completeness, resulting in 62% more variables with >95% completeness. Overall, mean variable completeness was 9.45% higher in Colo-661 following QC when compared with the original dataset (following missing value encoding, described above).

**Figure 4:**
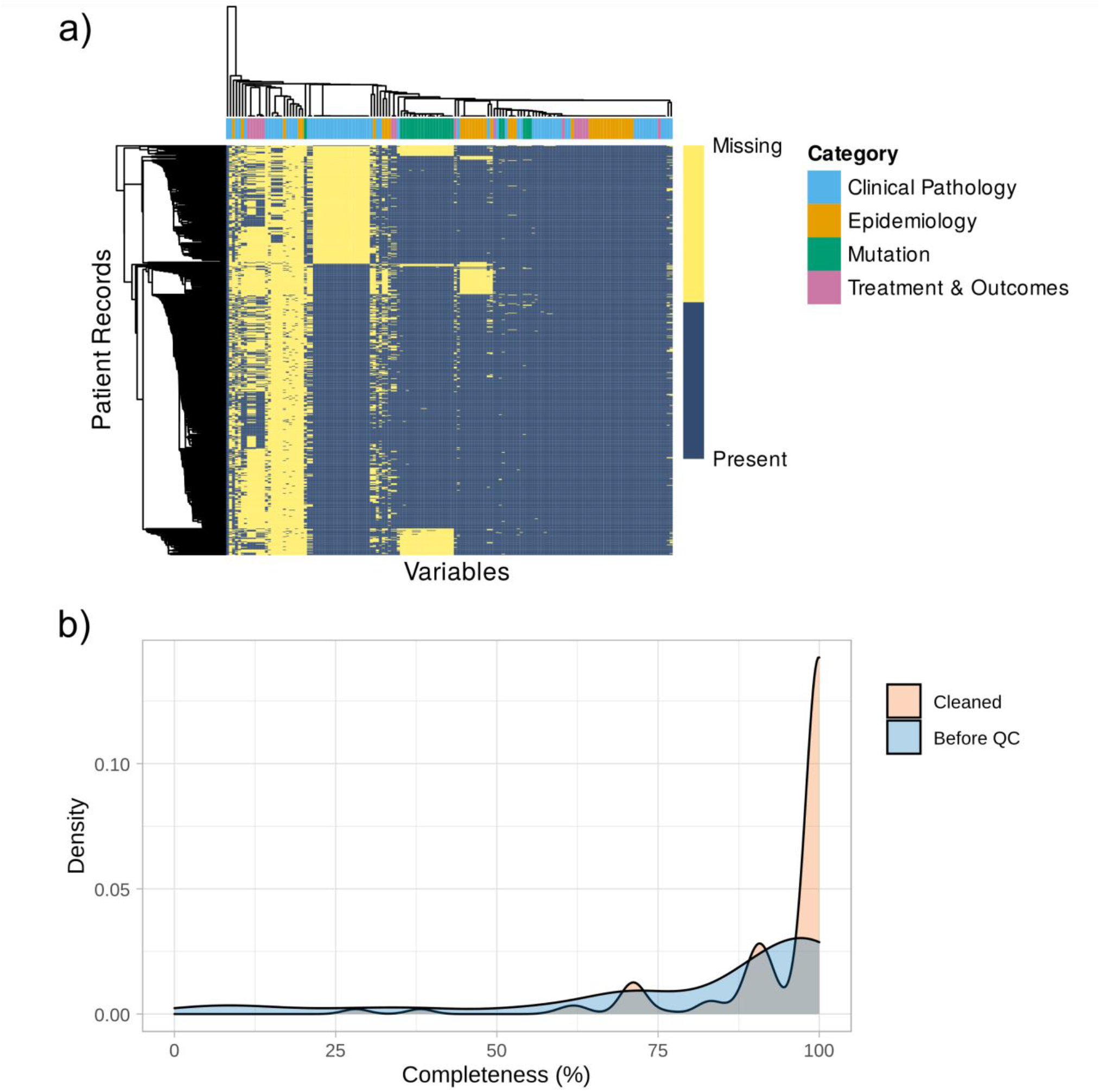
Characterising and comparative measures of completeness in Colo-661. a) Binary heatmap showing Colo-661 variable (x-axis) and patient record (y-axis) completeness. Non-numeric variables were numerically encoded by the number of unique values they possessed. Missing values were numerically encoded as a highly distant value. The dendrograms (left and top) reflect hierarchical clustering of the values’ Euclidean distances using single-linkage clustering agglomeration. Yellow cells represent missing values whereas blue cells represent present values. The user-defined data modalities are identified in the coloured bar along the top of the heatmap with a corresponding legend on the right. Multiple regularly shaped areas of missingness are visible and each block of missingness generally contains variables of the same type. b) A comparison of variable completeness in the dataset as received (blue) and the cleaned dataset (orange). QC improved overall completeness with a substantial increase in variables possessing >95% completeness. No variables in the cleaned dataset have <28% completeness.

**Figure 5:**
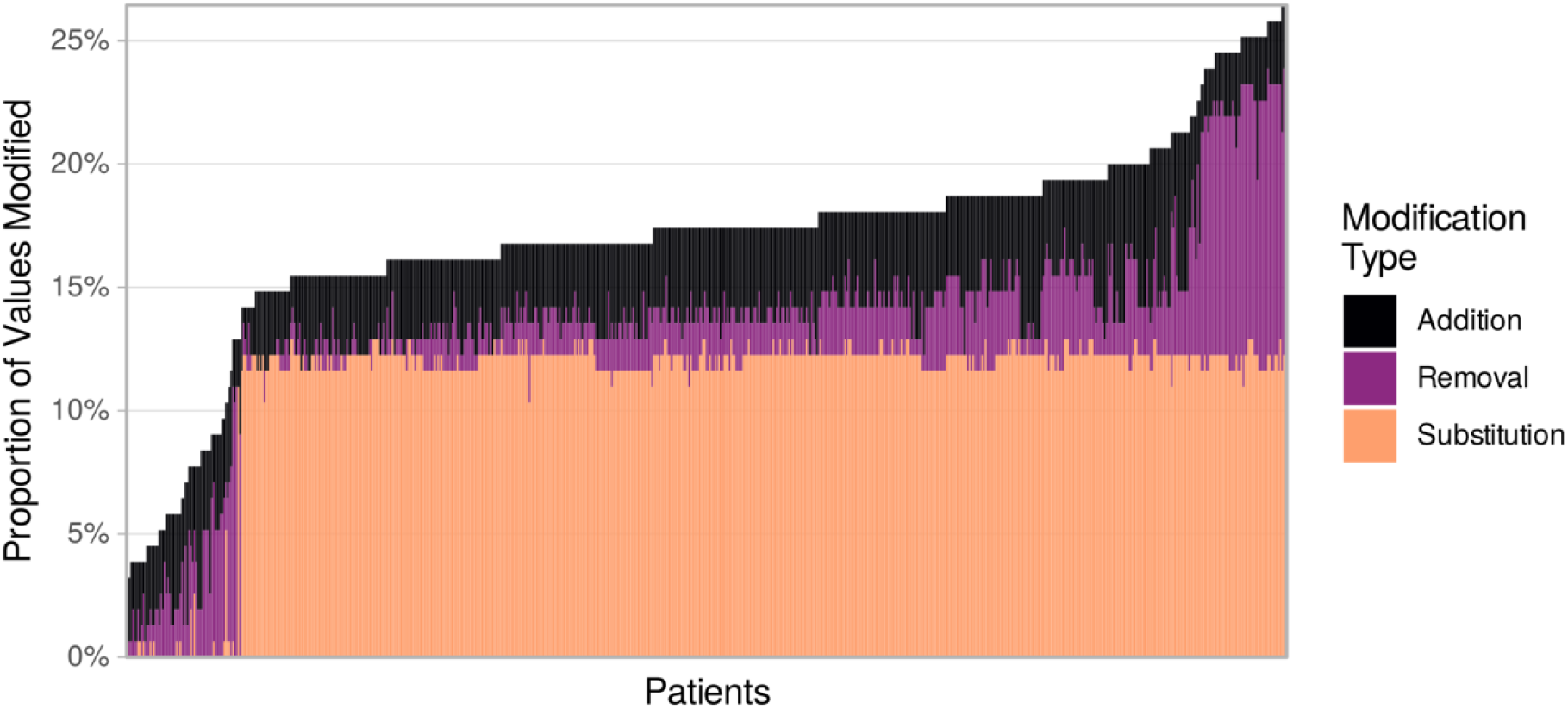
Uneven distribution of QC value modifications in the Colo-661 cohort. Stacked bar plot presenting modifications during QC as a percentage of total values per patient are shown on the y-axis. Patient records are displayed on the x-axis, ordered by y-axis values. The proportion of substitutions shows remarkable consistency across most patient records, due to the standardisation of mutation variables which altered all present values; although some patients had missing mutation values which could not be standardised and therefore do not appear.

### Semantic Enrichment

Ontologies contain valuable curated information about the relationships between domain concepts. Indeed, ontological relationships have been widely employed to support the interpretation of results from high-throughput technologies [34]. Rectangular health datasets (i.e., data frames or matrices) are semantically disorganised. However, ontologies can be utilised to capture semantic relationships between variables during data preparation in a process termed here as ‘semantic enrichment’. The structure of the ontology provides for aggregation values from constituent variables, in order to generate ‘meta-variables’. The process is explained below, as applied using the ‘semantic_enrichment’ function in eHDPrep, is summarised in Figure 6 and a worked example is provided (Supplementary Figure S8).

**Figure 6:**
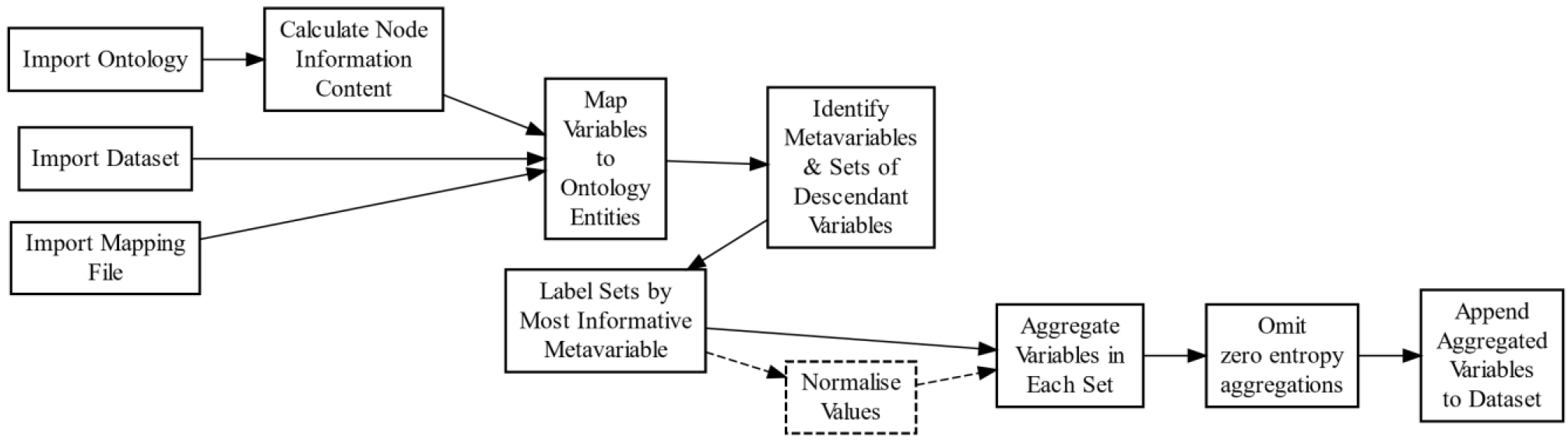
Semantic enrichment workflow in eHDPrep. The ordering of the steps shows logical dependencies. Dashed box and lines signify that ‘Normalise Values’ is an optional step, only required if variables have differing magnitudes. The ‘Map Variables to Ontology Entities’ step requires extensive user input. Meta-variables with zero entropy contain no information and so are omitted before the final step of appending meta-variables to the dataset.

### Discovery of the most informative common ancestor terms and semantic aggregation

The IC of each node in the supplied ontology is initially computed to quantify the specificity of nodes by depth and relative number of descendants [35]. Nodes representing variables, manually mapped to the ontology, are added to form an ontology:variable network. Sets of variables which share semantic commonalities through common ontological ancestors are identified. Sets of variables may have multiple common ancestors, therefore the IC of all common ancestors of a set are compared to identify the Most Informative Common Ancestor (MICA), which labels the set. Min-max normalisation (Equation (4) is applied to each variable prior to semantic aggregation whereby meta-variables are produced by taking the row-wise sum, minimum, maximum, average, and product of the set for each MICA. Only meta-variables with non-zero entropy (Equation (1) are appended to the dataset.

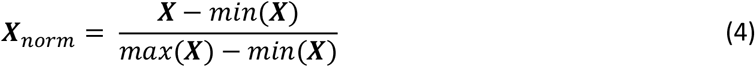

Where ***X*** is a numeric vector.

### Ontology Preparation

Variables in Colo-661 were mapped to two medical ontologies (Supplementary Table S1): the Systematized Nomenclature of Medicine Clinical Terms (SNOMED CT) ontology which standardises clinical terms for generation of electronic health records, covering >350,000 concepts at present [23]; and the Gene Ontology (GO), a widely used knowledgebase for gene function that currently contains >44,000 terms [36,37]. The Colo-661 variables were mapped to SNOMED CT by manual review assisted by the UK National Health Service Digital SNOMED CT Browser [38]. The UK edition of the SNOMED CT Clinical Edition ontology, version 31.1.0, was downloaded from the UK National Health Service’s technology reference data update distribution resource [39]. SNOMED CT was converted from Release Format 2 (RF2) to W3C Web Ontology Language (OWL) format using version 2.9.0 of the official SNOMED CT OWL toolkit [40]. We used ROBOT [41] to convert SNOMED from OWL to comma separated values containing each node’s superclasses, enabling generation of an edge table (Supplementary Figure S2) that defined the network graph to which the mapped Colo-661 variables were joined. For GO mapping, variables with gene assignments within the Colo-661 resource were verified and mapped to GO terms using Ensembl release 105 [42,43] via the BiomaRt package [44,45]. The ontologyIndex package [15] was used to import the January 2022 GO release into R. The Biological Process (BP) domain was chosen to create a network of Colo-661 variables and the mapped genes with their ‘is_a’ ontological ancestors.

### Enrichment Outcomes

A total of 193 (93.2%) of the post-QC Colo-661 variables were mapped to SNOMED CT or GO (Table 2, Supplementary Table S1). The remaining variables represented negative findings or missing values generated during one-hot encoding, which do not have equivalent entities in SNOMED CT (Supplementary Table S1) and would negate corresponding positive findings if mapped to their entities. For example, a finding of no diabetes mellitus (‘dm_type_NoDM’) or a missing entry for marital status (‘maritalcat_NA’). In total, 1600 meta-variables were generated and appended from 394 variable sets. Approximately seven times more variable sets were identified per mapped variable in the GO than in SNOMED CT with Colo-661; likely due to the 23.2 times greater mean number of direct annotations in GO which resulted in more common ancestors between variables (Table 2, Supplementary Table S1, Supplementary Table S3). The mean completeness of the Colo-661 meta-variables (98.7%) was 5.1% higher than their constituent variables.

**Table 2:** Summary statistics of ontologies applied to Colo-661 in semantic enrichment. The GO network has a higher mean number of direct annotations per mapped variable than SNOMED CT, which may explain the larger number of meta-variables generated using GO from fewer mapped variables. Of the 207 variables in the encoded Colo-661 dataset following completion of QC, 14 were not mapped to either SNOMED CT or GO. The number of nodes and edges were measured before the addition of mapped variable nodes.

| Source Ontology | Number of Nodes | Number of Edges | Number of Mapped Variables (Percentage of Total) | Mean Number of Direct Annotations per Mapped Variable | Number of Variable Sets | Number of Meta-variables |
| --- | --- | --- | --- | --- | --- | --- |
| SNOMED CT | 2448 | 4445 | 160 (77%) | 1.3 | 160 | 550 |
| Gene Ontology | 1675 | 2938 | 33 (16%) | 30.2 | 234 | 1050 |

The benefit of semantic enrichment is further demonstrated by 88.2% non-redundant information between the meta-variables from semantic aggregation and their constituent variables in Colo-661, measured by symmetric uncertainty (Equation (5) [46]. Despite low mean redundancy, nine meta-variables (0.6%) were fully redundant with a constituent variable. For example, for one MICA (‘beta adrenergic receptor blocking agent therapy’; SNOMED CT ID 439630003) two of its resultant meta-variables were fully redundant with one of the MICA’s two constituent variables, ‘bisoprolol_cat’ (Supplementary Table S4). The redundancy arose from the low number of constituent variables and the low number of unique values which could be taken (0 or 1) which inhibited creation of new information.

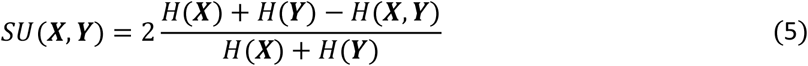

Where ***X*** and ***Y*** are numeric vectors and where H is entropy.

The SNOMED CT network, filtered to only include entities mapped to Colo-661 variables and their ancestors, was found to span many medical sub-domains. A selection of these domains are highlighted in Supplementary Figure S3 ranging from medical procedures, to substances, and to diseases. Selected SNOMED CT MICAs and their descendants are shown in Figure 7a and Supplementary Figures S4 and S5. The GO network was similarly filtered and connected several distinct biological concepts (Supplementary Figure S6) with example GO MICAs shown in Figure 7b and Supplementary Figure S7. In Figure 7a, five comorbidity variables are linked by their semantic commonality as types of heart disease while three mutation variables are linked by their involvement in drug catabolic process in Figure 7b. Figure 7a also visualises another MICA, ‘Ischaemic heart disease’, for two variables in the figure. Supplementary Figure S4 visualises the linkage of three variables, describing prescription medications and an adjuvant treatment regimen, as enzyme inhibitor products. The semantic commonalities between thirteen variables describing tumour excision location, other surgery types, and emergency surgery status are shown in Supplementary Figure S5 with the ‘Surgical procedure’ MICA. This figure contains additional MICAs for subsets of shown variables, for example node D (‘Right colectomy’). Supplementary Figure S7 shows commonality in a typical feature of several cancers, ‘Negative regulation of programmed cell death’ [47], between sixteen variables from multiple modalities.

**Figure 7:**
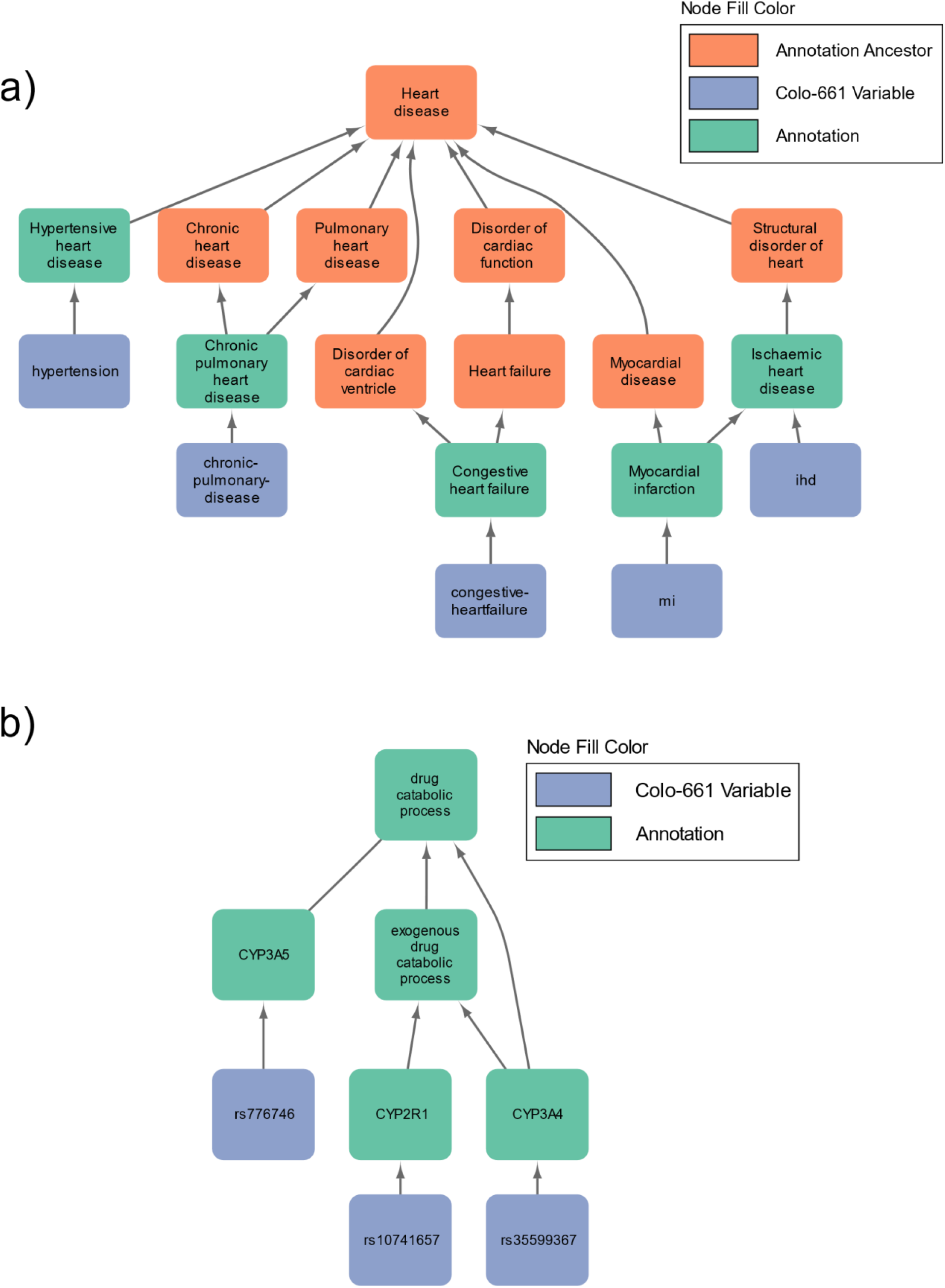
Exemplar Most Informative Common Ancestors (MICAs) and the semantic relationships with their constituent Colo-661 variables. a) Relationships between comorbidities are identified by semantic enrichment with SNOMED CT. ‘Heart disease’ is the MICA for all constituent variables (blue) shown in this figure however ‘Ischaemic heart disease’ is also a MICA for the Colo-661 variable set containing only the ‘mi’ and ‘ihd’ variables. b) Three single nucleotide polymorphism variables are semantically linked in the GO by the MICA ‘drug catabolic process’. Mutation variables in Colo-661 were selected based on prior association with colon cancer and exogenous exposures of interest. MICAs can help increase the interpretability of their constituent variables by integrating informative context encapsulated in the ontology; the resulting meta-variable may be useful in downstream analyses for example in feature selection for precision medicine applications.

## Methods

### Dataset

We utilised multi-modal data for a cohort of 661 patients as an exemplar for our approach, representing 89% of stage II/III colon adenocarcinoma patients who underwent surgery in two healthcare trusts in Northern Ireland between 2004 and 2008 (Colo-661). The cohort primarily comprised colon cancer patients, and two rectal cancer patients who did not undergo neo-adjuvant therapy. Follow-up concluded at the end of 2013 where 212 patients had died from a CRC-specific cause. Subsets of Colo-661 have been used in previous publications and the data had therefore undergone prior manual QC operations [19,20,48–54]. Researcher-defined data modalities span clinical pathology, epidemiology, mutation, and treatment and outcomes. These modalities originated from several sources: electronic health records, pathology reports, medical charts, the Northern Ireland Clinical Oncology Information System, Northern Ireland Registrar General’s Office, tumour image analysis [49], ColoCarta panel [55], and targeted mutation analysis.

### Data Processing

A combination of base R [56] and the Tidyverse family of packages [13] is used for data formatting and manipulation in eHDPrep. The quanteda package [12] is applied for parsing free text variables into computable units (tokens), token cleaning, and ultimately to extract information by identifying words which often appear near each other using quanteda’s ‘tokens_skipgrams’ function. The ‘stopwords’ function from tm [25,57] identifies stopwords, such as “a”, “in”, and “if”, for removal during cleaning. Cluster analysis is implemented through the dist and hclust functions in the stats package [56] to calculate Euclidean distance and single-linkage clusters, respectively, to analyse dataset completeness. Networks for semantic enrichment are created, manipulated, and analysed using igraph and Tidygraph [58,59].

### Data Reporting and Visualisation

The specificity and sensitivity of variables generated from free-text in Colo-661, following preprocessing, were assessed through comparative manual review between each value in the generated variables (n=3305) and corresponding value in the source variable. Two of the 3305 values were false positives (99.9% specificity) and 23 were false negatives (89.9% sensitivity). eHDPrep produces summary tables generated using the tibble package [60] which are optionally formatted using knitr [61,62] and kableExtra [63]. Heatmaps of dataset completeness are visualised with pheatmap [64] and the remaining plots are created with ggplot2 [65]. Network visualisations in this paper were generated using RCy3, Cytoscape, and InkScape [66–68]. Nodes in Supplementary Figures S3 and S5 were sized by PageRank centrality [69].

## Discussion

eHDPrep delivers an accessible set of functions, demonstrated here with suggested workflows applied to a real-world medical dataset (Colo-661), resulting in demonstrable benefit to data quality. Improvements to Colo-661 using eHDPrep included standardisation of eight strings representing missingness to “NA”, resolution of forty internal inconsistencies, conversion of free text to eleven new variables, numeric encoding of 123 nominal and ordinal variables, and a 9.45% increase in mean variable completeness. Additionally, we have showcased tools for assessment of data quality in both the input data and the data following QC operations.

eHDPrep provides novel functionality for data preparation in R where meta-variables are created by aggregation using ontological semantic commonalities. The benefit of this semantic enrichment is exemplified in Colo-661 through the creation of 1600 meta-variables. Furthermore, the mean redundancy of the meta-variables with their constituent variables was 11.8%, demonstrating creation of substantial information that was absent from the input dataset. The added non-redundant information in meta-variables may potentially enable discovery of patterns where the disaggregated data would be too heterogeneous or too sparse to identify meaningful results. We also observed a 5.1% higher average completeness of the meta-variables relative to their constituent variables. For patients where some values in constituent variables are missing, the meta-variables will contain semantic aggregations of the non-missing values, affording more comprehensive patient representation in downstream analyses while simultaneously preserving missing values that may be indicative of patient health and background [70,71]. A further benefit of eHDPrep is interoperability, an important consideration in digital healthcare [72,73]. The standardised encoding of the data improves syntactic interoperability and streamlines incorporation into larger databases while the meta-variables support semantic interoperability; for example, during data linkage in identifying similar variables across resources with differing degrees of data aggregation.

Semantic enrichment may be widely useful in health data analysis due to the availability of multiple rich, comprehensive ontologies; for example the Disease Ontology and the Human Phenotype Ontology [74,75]. The results of semantic enrichment in eHDPrep are critically dependent upon the ontology taken as input, and will likely suffer from a degree of annotation bias [76]. Also, mapping variables to ontology terms can be time-consuming and may require background knowledge of the variables if their labels are not self-explanatory; these issues may be mitigated by fuzzy string matching [77] and software interfaces, for example the UK National Health Service Digital SNOMED CT Browser [38]. Importantly, variables generated through semantic enrichment might not properly represent the quantitative relationship between their constituent variables due to unusual associations, such as J-curves [78]. Careful variable aggregation may be applied to avoid variation in two variables cancelling each other out. The identified semantic relationships may also aid in interpretation of why the variables have a particular association. For example, if opposite J-curves were found for values of the variables in Figure 7b, their common involvement in drug catabolism might help to understand the pattern of association.

The improved data quality, interoperability, and meta-variables generated through semantic enrichment in eHDPrep is expected to provide for greater robustness and added value in downstream analyses of biomedical data, including Colo-661.

## User documentation and technical details

eHDPrep contains short-form documentation for each function; called with ?[function name]. Long-form documentation, known as a vignette, is also provided to demonstrate QC and semantic enrichment functionality with synthetic example data, R code, and explanatory text. The vignette is created when the package is built and reflects the functionality of the current version. Error and warning handling messages have been included to ensure expected inputs are received and to notify if unexpected outcomes are returned. eHDPrep is written in the R programming language with a codebase size of 3943 lines of code and 57 unit tests.

## Availability of supporting source code and requirements

Project name: eHDPrep

Project home page: https://github.com/overton-group/eHDPrep. [[CRAN link will go here]].

Operating systems: Windows and Linux

Programming language: R

Other requirements: R (≥ 3.6.3)

Licence: GPLv3

## Availability of supporting data

Access to the Colo-661 dataset may be requested by contacting the Northern Ireland Biobank [79] (; director Prof. J. James). Synthetic demonstrator data are available within the eHDPrep package.

## List of abbreviations

CRC: colorectal cancer
QC: quality control
GO: gene ontology
SNOMED CT: systematized nomenclature of medicine clinical terms
IC: information content
MIC: mutual information content
MICA: most informative common ancestor.

## Ethics approval and consent to participate

Provision and use of this dataset was approved by the Epi700 Consortium and Northern Ireland Biobank under Secondary Use of Data. REC references: 10/NIR02/53, 11/NI/0013.

## Consent for publication

Not applicable.

## Authors’ contributions

Conceptualization, IO; Data curation, TT, HC, IO; Formal analysis, TT, IO; Funding acquisition, PM, HC, IO; Interpretation TT, TF, HC, IO; Investigation, TT, IO; Methodology, TT, HC, IO; Project administration, IO; Resources IO; Software TT, IO; Supervision, PM, TF, HC, IO; Validation TT, HC, IO; Visualization, TT, PM, HC, IO; Writing—original draft, TT, IO; Writing—review and editing, TT, PM, TF, HC, IO. All authors have read and agreed to the published version of the manuscript.

## Acknowledgements

We are grateful to the Epi700 Steering Committee for curation of and support with the data resource containing Colo-661. The samples used in this research were received from the Northern Ireland Biobank which has received support from Health and Social Care Research and Development Division of the Public Health Agency in Northern Ireland and Cancer Research UK (via the former Belfast CRUK Centre and the Northern Ireland Experimental Cancer Medicine Centre) and the Friends of the Cancer Centre. The Northern Ireland Molecular Pathology Laboratory, which was responsible for creating resources for the Northern Ireland Biobank, has received funding from Cancer Research UK, the Friends of the Cancer Centre and the Sean Crummey Foundation. Clinical data collected and analysed was facilitated by The Northern Ireland Cancer Registry, which is funded by the Public Health Agency, Northern Ireland. Thanks to Dr. Alexander Lubbock and Dr. Seanna McTaggart for helpful discussions, also to the Overton group for testing eHDPrep.

## Authors’ information

TT is a PhD candidate in the Overton group, Patrick G Johnston Centre for Cancer Research, Queen’s University Belfast (QUB). PM is a Reader, theme lead for Cybersecurity and deputy director of The Centre for Secure Information Technologies, QUB. TF is team lead for Data Science at LifeArc’s Centre for Diagnostics Development. HC is Professor of Cancer Epidemiology and the Cancer Epidemiology research group lead in the Centre for Public Health and Patrick G Johnston Centre for Cancer Research, QUB. IO is a Senior Lecturer in Data Intensive Biomedicine and Medical Bioinformatics Research Section head in the Patrick G Johnston Center for Cancer Research, QUB.

## Supplementary Material

**Supplementary Figure S1:**
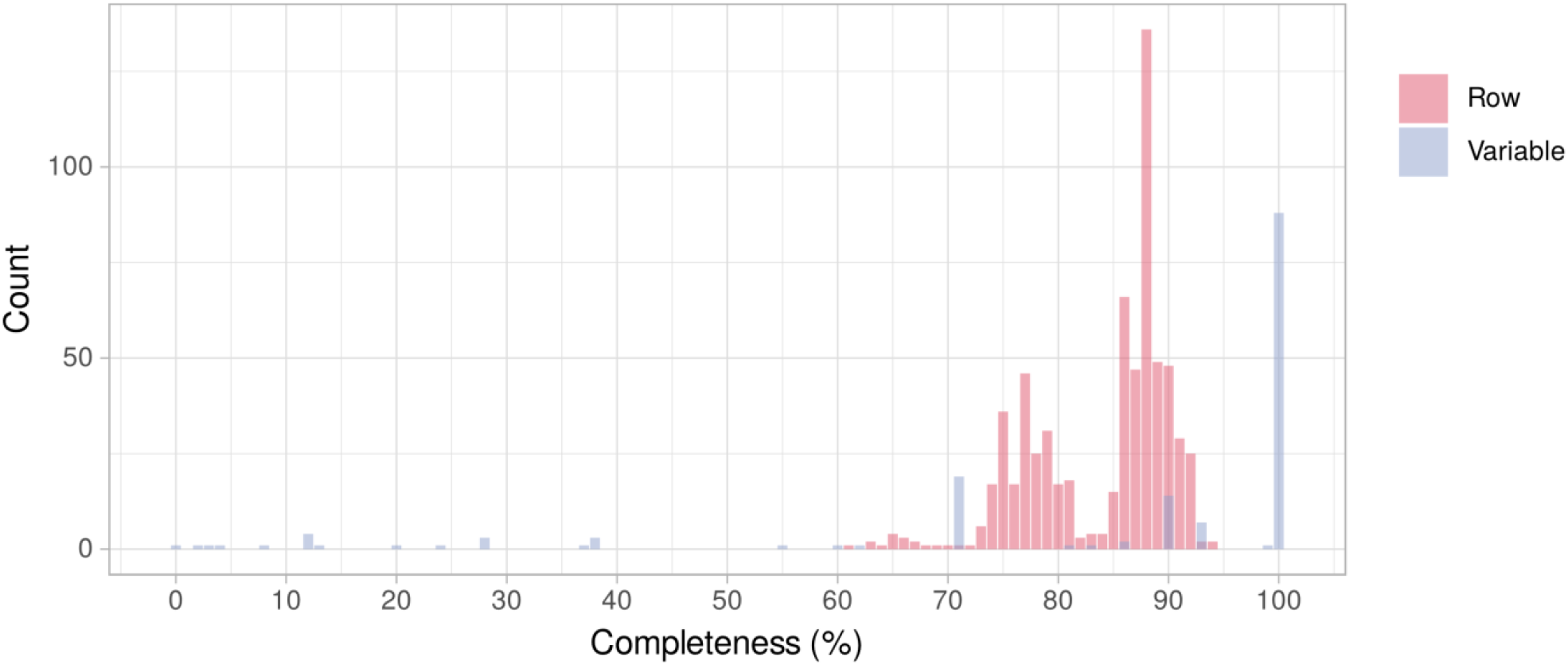
Completeness in Colo-661. This bar plot summarises patient record (red) and variable (lilac) completeness in the unprocessed Colo-661 dataset. Patient records (red) were between 61% and 94% complete while variable completeness (blue) ranged from 0% to 100%.

**Supplementary Figure S2:**
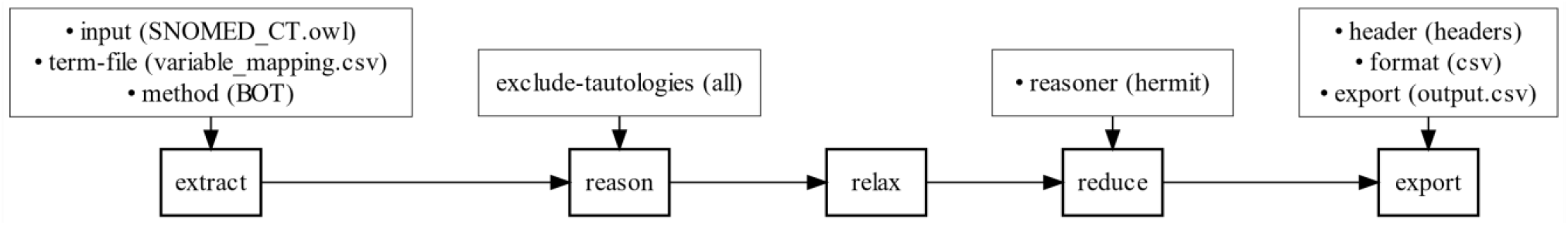
Preparation of SNOMED CT for semantic enrichment. ROBOT commands are joined with horizontal arrows. The arguments are shown above their corresponding commands, joined with a vertical arrow with applied parameters displayed in brackets. The “extract” command was used to subset the input ontology. The “BOT” method subset all terms in the ontology to entities mapped to Colo-661 variables plus all super-classes and inter-relations between super-classes. The “reason” command was used to logically validate and automatically classify the ontology using the reasoner “hermit” with all tautologies removed. The “relax” command was used to relax Equivalence axioms to weaker SubClassOf axioms which is suitable for semantic enrichment. The “reduce” command removed redundant SubClassOf axioms using the “hermit” reasoner. Finally, the “export” command exported the ontology as a comma separated values for import into R.

**Supplementary Figure S3:**
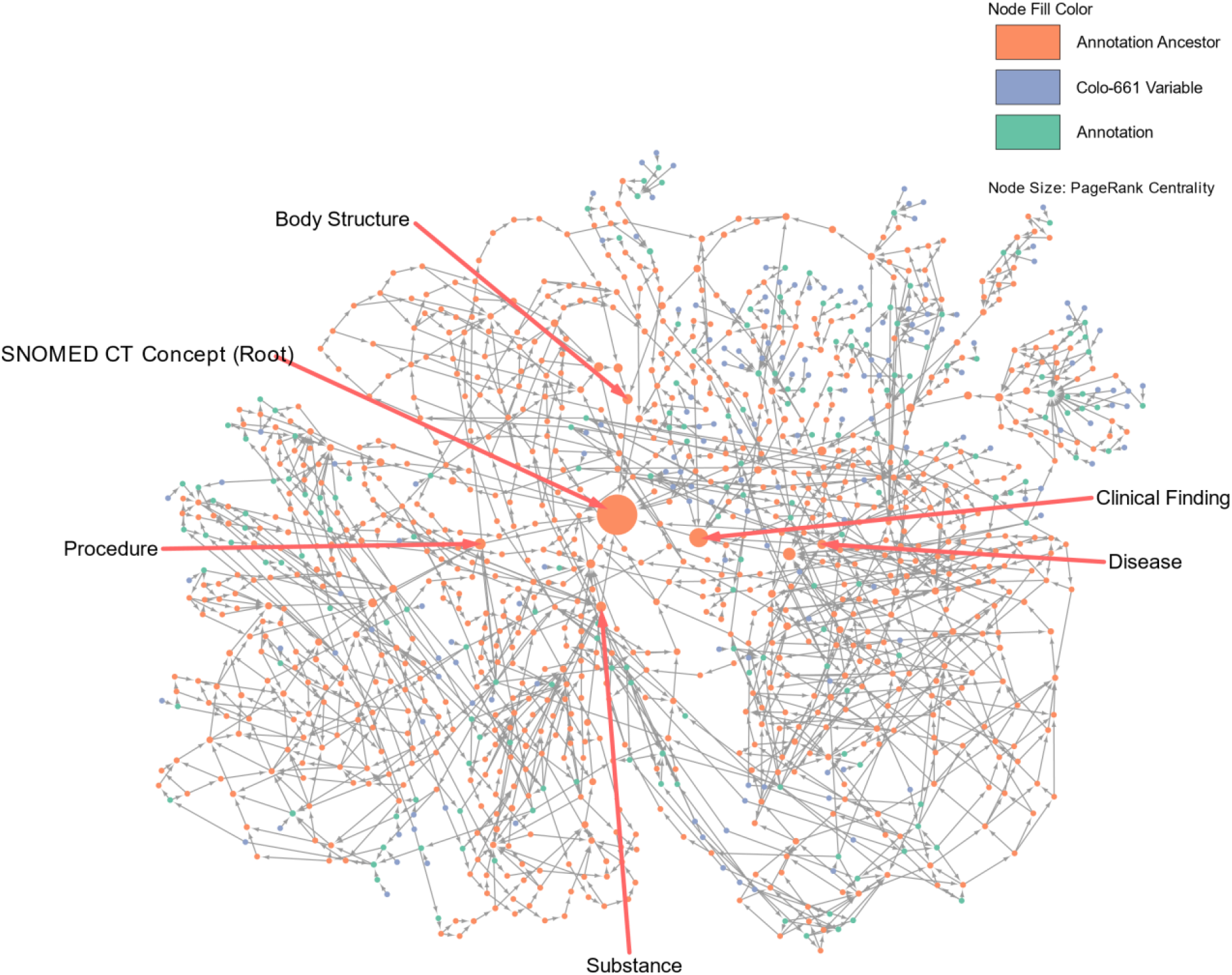
SNOMED CT annotation network. Network nodes represent Colo-661 variables (blue), SNOMED CT terms mapped to Colo-661 variables (green) and their ancestor ontology terms (orange). Node size is proportional to PageRank centrality [69]. The large nodes are highly central, representing domains within the network, some of these are labelled with their SNOMED CT term names (red arrows).

**Supplementary Figure S4:**
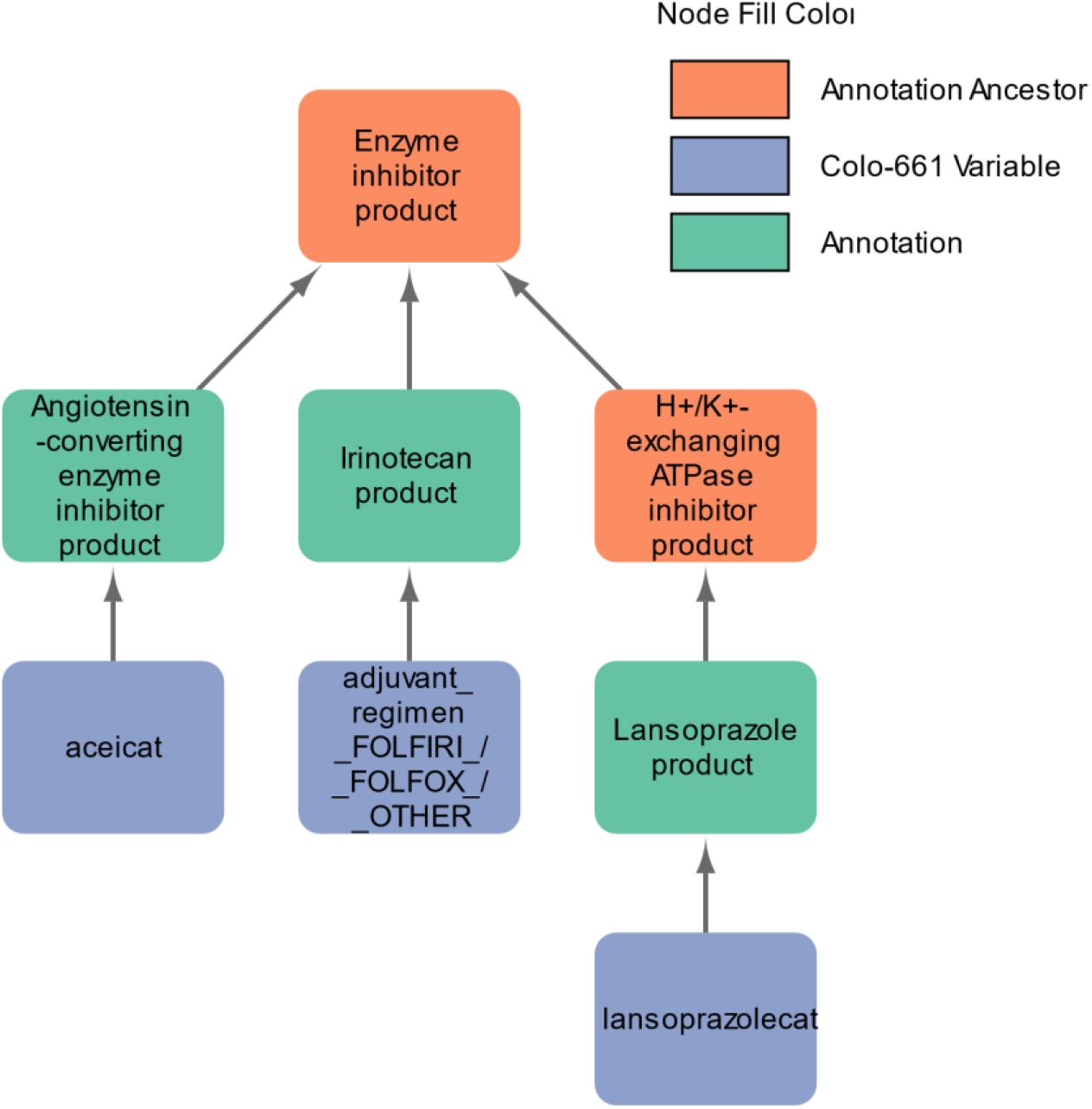
Example of a Most Informative Common Ancestor (MICA) term ‘Enzyme inhibitor product’. Variables describing two medications (‘aceicat’ and ‘lansoprazolecat’) and an adjuvant regimen were semantically linked through the SNOMED CT term ‘Enzyme inhibitor product’. Therefore, semantic enrichment of Colo-661 identified a degree of functional similarity across distinct treatment regimes.

**Supplementary Figure S5:**
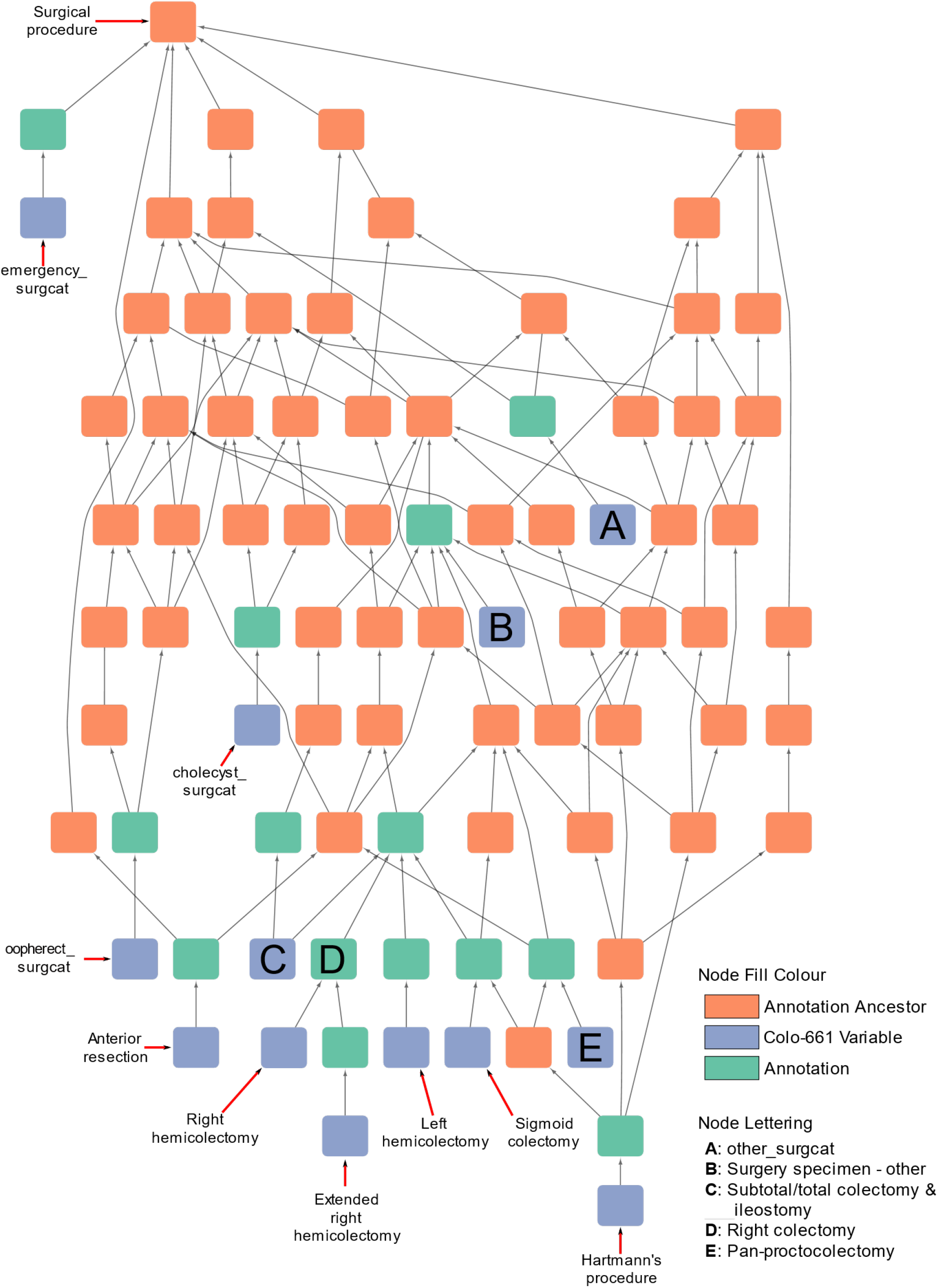
SNOMED CT Network for Most Informative Common Ancestor (MICA) term ‘Surgical procedure’. Colo-661 variables describing the procedure used to excise the primary tumour, or describing other operations, and the emergency status of the patient’s operation are semantically linked via the ‘Surgical procedure’ MICA. This MICA encompasses a relatively large number of variables (n=13) aggregating information across a range of surgical procedures that could be useful in later analyses. This network also includes several other MICAs, corresponding to smaller groupings of Colo-661 variables, such as node D (Right colectomy), which aggregates the variables ‘Right hemicolectomy’ and ‘Extended right hemicolectomy’.

**Supplementary Figure S6:**
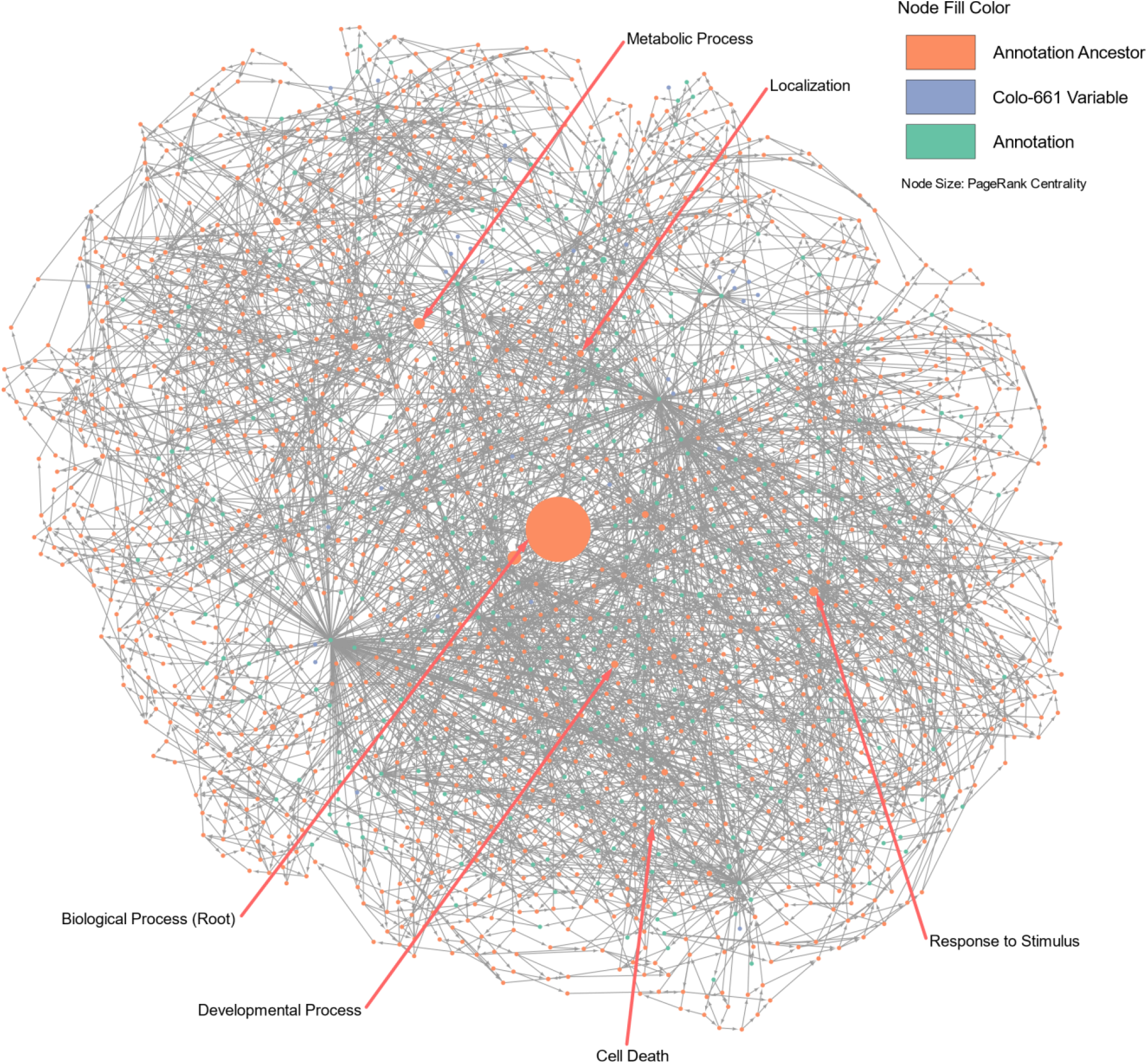
The Colo-661 Gene Ontology (GO) Biological Process annotation network. Network nodes show Colo-661 variables (blue), their mapped genes (green) with associated GO terms (green), and ancestor terms (orange). Node size is proportional to PageRank centrality [69]. Larger nodes have high PageRank centrality, represent domains within the network and some of these are labelled (red arrows) with their names from GO.

**Supplementary Figure S7:**
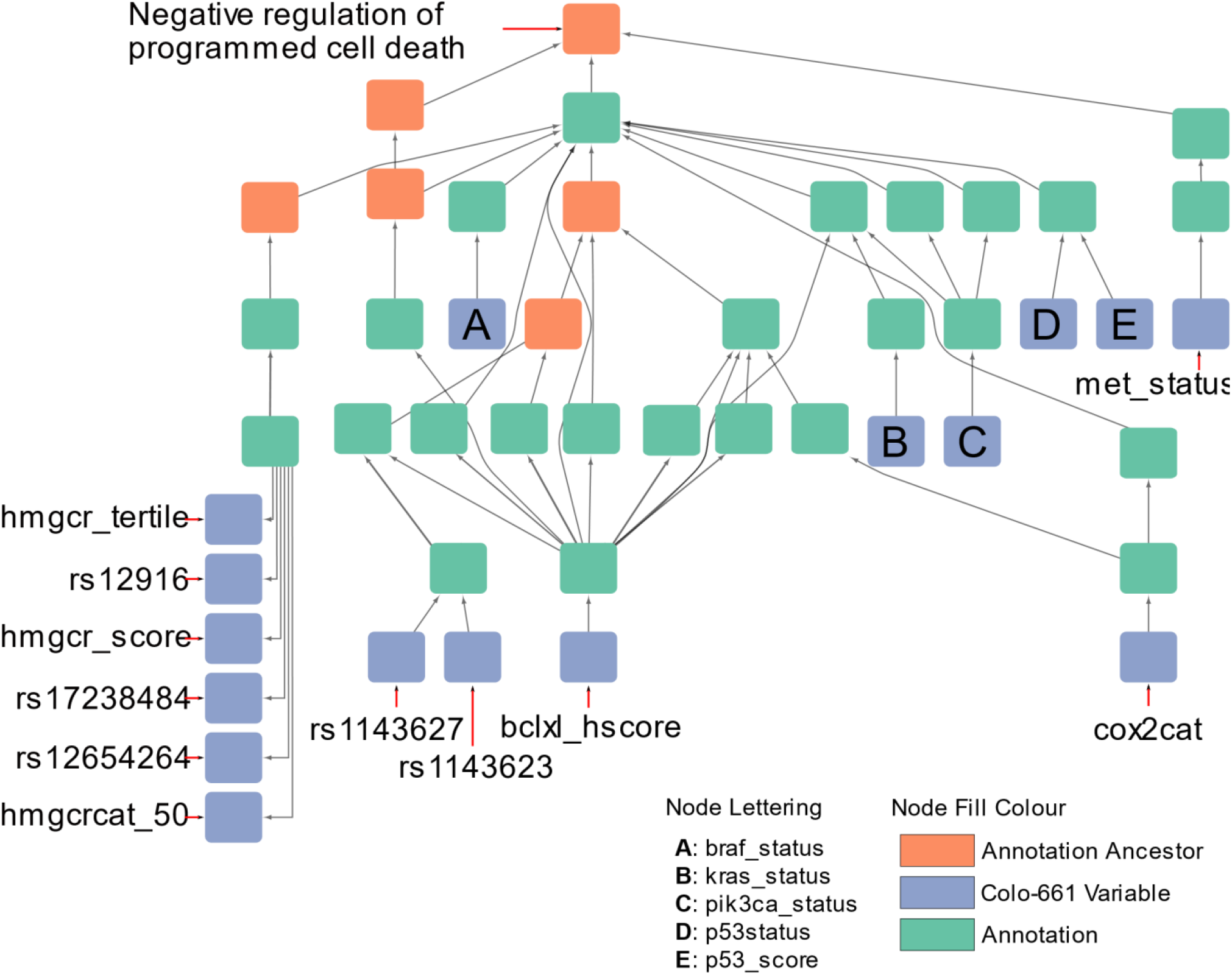
Gene Ontology Biological Process Network for the Most Informative Common Ancestor (MICA) term ‘Negative regulation of programmed cell death’. Network nodes show Colo-661 variables (blue), their mapped genes (green) with associated GO terms (green), and their ancestor terms (orange). The ‘Negative regulation of programmed cell death’ MICA describes an important step in the progression of many cancers [47], where cells can evade signals that lead to cell death. Additionally, the Figure exemplifies aggregation of variables from different data modalities. For example, ‘hmgcr_tertile’ and ‘rs12916’ at the bottom left of the figure are immunohistochemical and single nucleotide polymorphism variables, respectively.

**Supplementary Figure S8:**
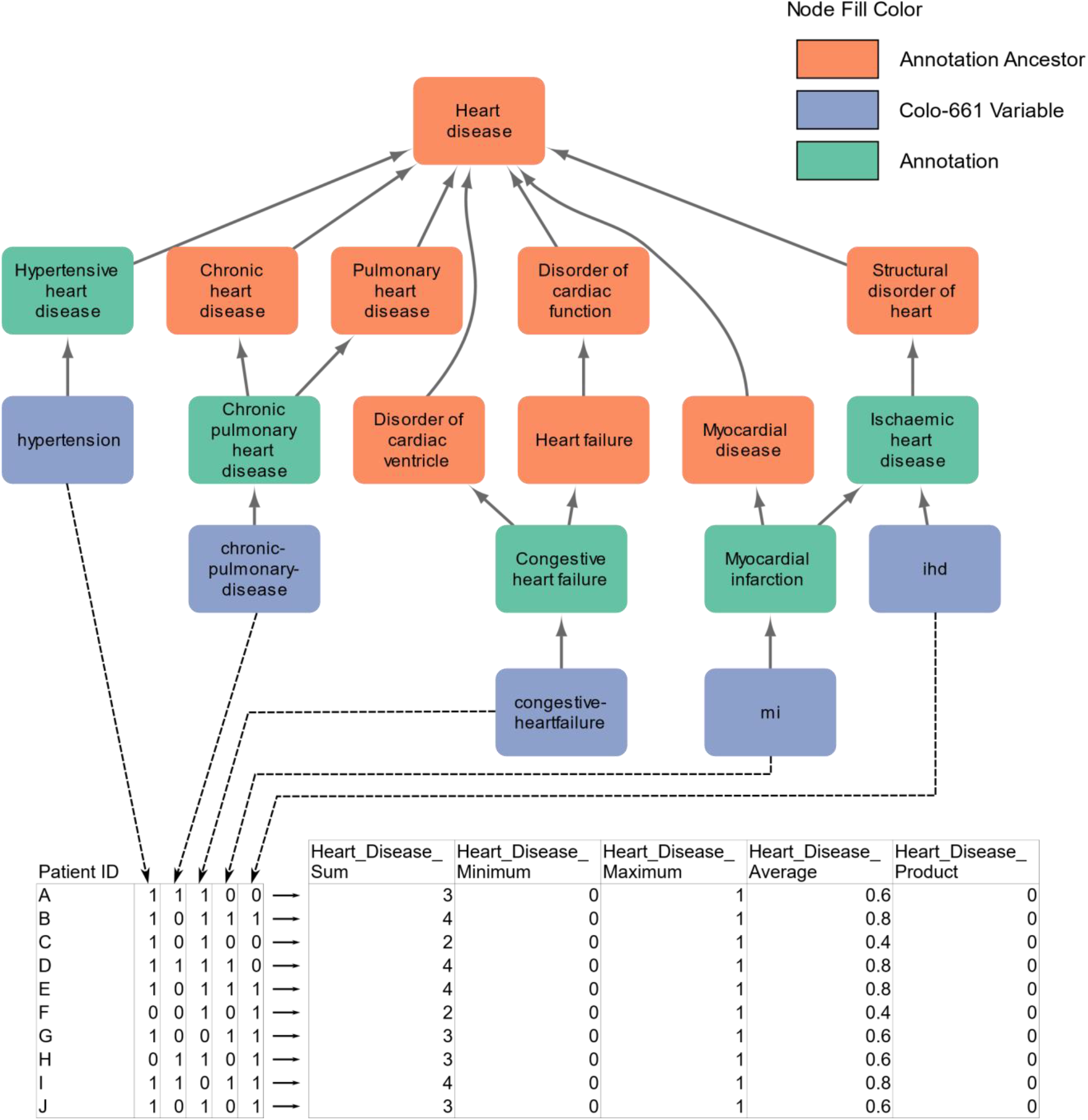
Worked example of aggregation in semantic enrichment. Five variables from Colo-661 in the network (blue) were mapped to entities in SNOMED CT that share ‘Heart disease’ as their Most Informative Common Ancestor (MICA). These five variables therefore constitute a “set” and are selected from example synthetic data containing these variables’ names (table, bottom-left) and are aggregated row-wise. The aggregated variables (table, bottom-right) are appended to the synthetic dataset, labelled with the MICA’s name and the corresponding aggregation function.

**Supplementary Table S1: List of Colo-661 variables and ontological mappings. [Provided as a separate file]**. Variables which were added, preserved or removed can be identified by the “Presence Post-QC” column. The user-defined variable modality is recorded in the “Modality” column. Data classes, as encoded in R, are given in the “Data Types” columns. Ranges of values are shown in “Value Range in Post-QC Dataset”; no range is given if a variable was removed from the dataset during QC. Our mapping(s) are provided in the columns “Mapped Ontology” and “Ontological / Gene Mapping”. Variables mapped to the GO require an initial mapping to a gene, shown here, with Gene:GO term mappings detailed in Supplementary Table S3.

**Supplementary Table S2: Internal consistency checks performed on Colo-661. [Provided as a separate file]**. Tests between variables both containing numeric values were performed using the logical operator in ‘Logical Test’ with the format ‘[Variable A] [Operator] [Variable B]’. Between variables containing categories, values in ‘Variable B Boundaries’ were tested to only be present given the corresponding values in ‘Variable A Boundaries’. Tests between numeric and categorical values were similarly compared with numeric (inclusive) ranges denoted by colon-separated values.

**Supplementary Table S3: List of genes mapped to Colo-661 variables and the mapped GO terms. [Provided as a separate file]**. GO terms are separated by “;”. Mappings between genes and GO terms were sourced from Ensembl.

**Supplementary Table S4:**
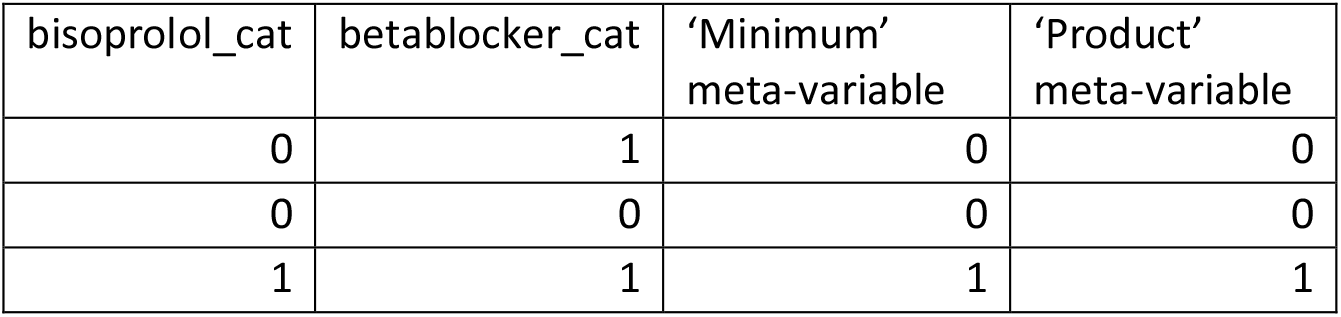
Redundancy between meta-variables and a consituent variable. Two of five meta-variables derived from minimum and product aggregations of ‘bisoprolol_cat’ and ‘betablocker_cat’ (semantically linked by the MICA: ‘beta adrenergic receptor blocking agent therapy’) were fully redundant with ‘bisoprolol_cat’. The table describes the observed row-wise combinations of values across the constituent variables and the two meta-variables which were redundant with ‘bisoprolol_cat’. While the value of ‘betablocker_cat’ differed from values of the meta-variables shown, the value of ‘bisoprolol_cat’ did not which led to the observed redundancy. ‘bisoprolol_cat’ and ‘betablocker_cat’ described if patients were prescribed bisoprolol and beta blockers, respectively.

| bisoprolol_cat | betablocker_cat | ‘Minimum’ meta-variable | ‘Product’ meta-variable |
| --- | --- | --- | --- |
| 0 | 1 | 0 | 0 |
| 0 | 0 | 0 | 0 |
| 1 | 1 | 1 | 1 |

